# Mechanical Property Based Brain Age Prediction using Convolutional Neural Networks

**DOI:** 10.1101/2023.02.12.528186

**Authors:** Rebecca G. Clements, Claudio Cesar Claros-Olivares, Grace McIlvain, Austin J. Brockmeier, Curtis L. Johnson

## Abstract

Brain age is a quantitative estimate to explain an individual’s structural and functional brain measurements relative to the overall population and is particularly valuable in describing differences related to developmental or neurodegenerative pathology. Accurately inferring brain age from brain imaging data requires sophisticated models that capture the underlying age-related brain changes. Magnetic resonance elastography (MRE) is a phase contrast MRI technology that uses external palpations to measure brain mechanical properties. Mechanical property measures of viscoelastic shear stiffness and damping ratio have been found to change across the entire life span and to reflect brain health due to neurodegenerative diseases and even individual differences in cognitive function. Here we develop and train a multi-modal 3D convolutional neural network (CNN) to model the relationship between age and whole brain mechanical properties. After training, the network maps the measurements and other inputs to a brain age prediction. We found high performance using the 3D maps of various mechanical properties to predict brain age. Stiffness maps alone were able to predict ages of the test group subjects with a mean absolute error (MAE) of 3.76 years, which is comparable to single inputs of damping ratio (MAE: 3.82) and outperforms single input of volume (MAE: 4.60). Combining stiffness and volume in a multimodal approach performed the best, with an MAE of 3.60 years, whereas including damping ratio worsened model performance. Our results reflect previous MRE literature that had demonstrated that stiffness is more strongly related to chronological age than damping ratio. This machine learning model provides the first prediction of brain age from brain biomechanical data—an advancement towards sensitively describing brain integrity differences in individuals with neuropathology.

## 1. Introduction

Throughout life, the brain changes in structural integrity, functional connectivity, tissue anisotropy, and synaptic plasticity, with brain degeneration rapidly accelerating in older adults (Betzel et al., 2014; Freitas et al., 2011; Takao et al., 2012). Understanding the natural progression of brain changes is critical toward establishing brain biomarkers of atypical development and degeneration. Advanced neuroimaging techniques capable of measuring structural integrity of neural tissue offer sensitive metrics that reflect glial matrix integrity, neuronal density, and myelin integrity (Hiscox et al., 2021; Johnson & Telzer, 2018). Magnetic resonance elastography (MRE) noninvasively images brain mechanical properties *in vivo*, which are sensitive to degeneration in the brain caused by aging and disease (Murphy et al., 2019). MRE has shown that the global brain softens because of aging at a rate of around 0.3-1% per year, with the specific softening rate varying among brain regions (Arani et al., 2015; Delgorio et al., 2021; Sack et al., 2011). The age-related softening of the brain can be exacerbated by neurodegenerative disorders such as Alzheimer’s disease, multiple sclerosis, and Parkinson’s disease (Delgorio et al., 2023; Hiscox, Johnson, McGarry, et al., 2020; Lipp et al., 2013; Murphy et al., 2011; Wuerfel et al., 2010). The sensitivity of mechanical property measures from MRE to tissue integrity make them strong candidates for imaging metrics used to predict brain health outcomes.

Machine learning provides an avenue for identifying patterns from dense, high-dimensional data to predict outcomes of interest. Specifically, convolutional neural networks (CNNs) extract both regional patterns and high-level structural patterns from 3-dimensional imaging data. CNNs learn to extract key features from images through a series of convolutional filters, pooling layers, and non-linear functions. Finally, the set of image features are fed into classification layers which relate features to subject characteristics through linear or non-linear regression. This is a supervised learning process wherein the true outcome is provided, and the weights of the regression equations are updated to minimize the error between target and predicted output. After training, given a new input without a target, the model can utilize the learned regression values to predict the output. CNNs have been used for several 3D medical imaging applications including tumor detection, disease progression prediction, image segmentation, and image reconstruction (El-Sappagh et al., 2020; Gupta et al., 2018; Moeskops et al., 2016; Rehman et al., 2021).

The use of CNNs to predict age from neuroimaging data has emerged as a popular area of research. By identifying features from imaging data that best describe age, one can use the CNN model to determine deviations between chronological age and expected age from images – “brain age” – to characterize brain health. Imaging modalities including diffusion, functional, and structural MRI have already shown promising results for brain age prediction (Chen et al., 2020; Cole et al., 2017; Jiang et al., 2020; Lund et al., 2022). The incorporation of MRE data into brain age models through machine learning offers a potentially powerful additional predictor to determine brain age given the high sensitivity of mechanical properties to age and disease. While aging effects on brain stiffness are well-studied (Hiscox et al., 2021), previous works have focused on region-of-interest (ROI) based analyses (Arani et al., 2015; Delgorio et al., 2021; Hiscox et al., 2018; Kalra et al., 2019; Takamura et al., 2020) and have not taken advantage of voxel-wise mechanical property measures from MRE (Hiscox, McGarry, et al., 2020).

Here, we aim to develop a 3D CNN that can learn age from whole brain maps of structural integrity measures including from MRE. We produce a network that can synthesize multiple 3D input maps for each subject as well as other demographic attributes as categorical variable inputs. Given that brain MRE is not widely used and published sample sizes are relatively small, we pooled data for this study from multiple studies employing similar imaging and processing protocols, as has been previously done by Hiscox et al. (Hiscox, McGarry, et al., 2020) We include data from healthy subjects across the life span (Figure 1) to allow us to model brain age with a reference of chronological age to ultimately provide a framework for understanding brain age in individuals with pathology. We demonstrate the high performance of MRE measures alone in predicting chronological age, use Bayesian optimization to determine the best CNN model hyperparameters, and explore model performance when combining MRE measures of brain stiffness with other imaging measures.

**Figure 1.**
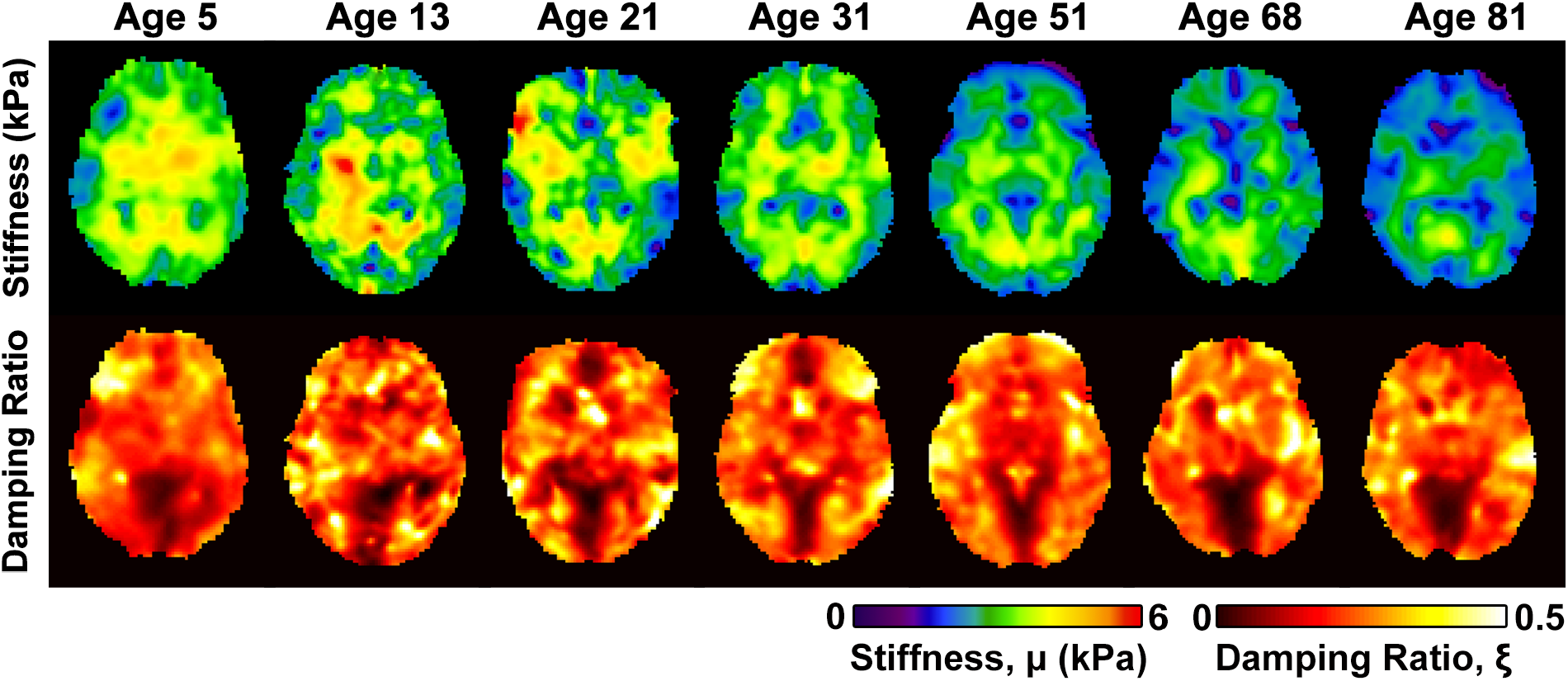
Example stiffness and damping ratio maps for cognitively normal subjects of varying ages.

## 2. Methods

### 2.1. Participants

The total dataset comprised 279 subjects ages 5–81 years (116 males, 163 females), with a mean age of 31.0 ± 21.3 years. This data was compiled from 8 studies with highly similar MRI protocols, which included an MRE scan and a high resolution T1-weighted anatomical scan, collected at the University of Delaware (UD) with a Siemens 3T Prisma MRI scanner or at the University of Illinois at Urbana-Champaign (UIUC) with a Siemens 3T Trio MRI scanner. An overview of the participant demographics and imaging parameters for each study is shown in Table 1. Informed written consent was provided prior to scanning by all study participants and guardians of study participants under the age of 18, and all study protocols were approved by the institutional review boards of the respective institutions. Data collected at UIUC was previously pooled and used to create a standard-space atlas of brain mechanical properties (Hiscox, McGarry, et al., 2020). Additional data included in this work has been reported, in part, in several previous publications (Delgorio et al., 2022, 2023; McIlvain, Schneider, et al., 2022; Sanjana et al., 2021; Schneider et al., 2022). Readers are referred to prior works for additional information.

**Table 1.** Overview of the demographics and imaging parameters for each study

| Study | 1 | 2 | 3 | 4 | 5 | 6 | 7 | 8 |
| --- | --- | --- | --- | --- | --- | --- | --- | --- |
| **N** | 9 | 30 | 61 | 19 | 23 | 54 | 76 | 10 |
| **# M/F** | 1/8 | 1/29 | 27/24 | 12/7 | 15/8 | 24/30 | 34/42 | 4/6 |
| **Age Range** | 20-60 | 18-32 | 18-35 | 12-15 | 5-8 | 14-73 | 23-81 | 12-14 |
| **Site** | UIUC | UIUC | UIUC | UD | UD | UD | UD | UD |
| **Scanner Model** | Trio | Trio | Trio | Prisma | Prisma | Prisma | Prisma | Prisma |
| **Number of coils** | 32 | 32 | 32 | 64 | 64 | 64 | 64 | 64 |
| **T<sub>1</sub> MPRAGE** |  |  |  |  |  |  |  |  |
| **TE (ms)** | 2.32 | 2.32 | 2.32 | 2.32 | 2.32 | 2.32 | 2.32 | 2.32 |
| **TR (ms)** | 1900 | 1900 | 1900 | 2300 | 2300 | 2300 | 2300 | 2300 |
| **Resolution (mm<sup>3</sup>)** | 0.9 | 0.9 | 0.9 | 0.9 | 0.9 | 0.9 | 0.9 | 0.9 |
| **MRE** |  |  |  |  |  |  |  |  |
| **Frequency (Hz)** | 50 Hz | 50 Hz | 50 Hz | 50 Hz | 50 Hz | 50 Hz | 50 Hz | 50 Hz |
| **Sequence** | Spiral | Spiral | Spiral | Spiral | Spiral | Spiral | Spiral | Spiral |
| **Resolution (mm<sup>3</sup>)** | 1.6 | 2.0 | 1.6 | 1.5 | 1.5 | 1.5 | 1.25 | 1.5 |
| **Number of slices** | 60 | 60 | 60 | 80 | 64 | 80 | 96 | 80 |

### 2.2. Imaging Data and Pre-Processing

All imaging data included a whole-brain, high-resolution T1-weighted magnetization-prepared rapid acquisition gradient echo (MPRAGE) with 0.9 mm^3^ isotropic resolution and a high-resolution MRE scan. MRE data acquisitions were taken using either the 3D multislab, multishot spiral sequence (Johnson et al., 2014) or the 3D multiband, multishot spiral sequence (Johnson et al., 2016; McIlvain, Cerjanic, et al., 2022), with resolutions ranging from 1.25 mm to 2 mm isotropic and whole-brain coverage. Imag<u>es were</u> downsampled to 2 mm resolution prior to analysis. All MRE data was collected using a pneumatic actuator pillow system (Resoundant, Rochester, MN) using 50 Hz vibration. Each MRE scan had a corresponding field map of the same resolution and brain coverage, which was used during image reconstruction to minimize geometric distortion (McIlvain et al., 2022).

Displacement images from MRE were converted into maps of mechanical properties using the nonlinear inversion (NLI) algorithm (McGarry et al., 2012), which returns estimates of the complex-valued shear modulus, *G*^*\**^ = *G*’+ *iG*”, where *G*’is the storage modulus and *G*” is the loss modulus. Mechanical property measures of stiffness (*μ*) and damping ratio (*ξ*) are calculated as 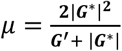 and *ξ* = *G*″/(2*G*^′^) (Manduca et al., 2001; McGarry & van Houten, 2008).

MRE magnitude data and the high resolution MPRAGE data were skull stripped using the Brain Extraction Tool (BET) in the FMRIB Software Library (FSL) (Jenkinson et al., 2012). The MRE images were registered to the corresponding T1-weighted anatomical image using FSL FLIRT (Jenkinson et al., 2002), which allows for an affine linear transformation, and the transform was similarly applied to the individual mechanical property maps. The T1-weighted anatomical scan was registered to the standard space MNI152 2-mm template with dimensions 91 × 109 × 91. Registration was done using a nonlinear registration with FSL FNIRT (Andersson et al., 2007) and the nonlinear transform was applied to the lower resolution registered MRE mechanical property maps. The result was all stiffness and damping ratio maps in identical spatial locations but retaining their voxel level stiffness values. Example anatomical scans and maps of stiffness and damping ratio before and after conversion to standard space are shown in Figure 2A.

**Figure 2.**
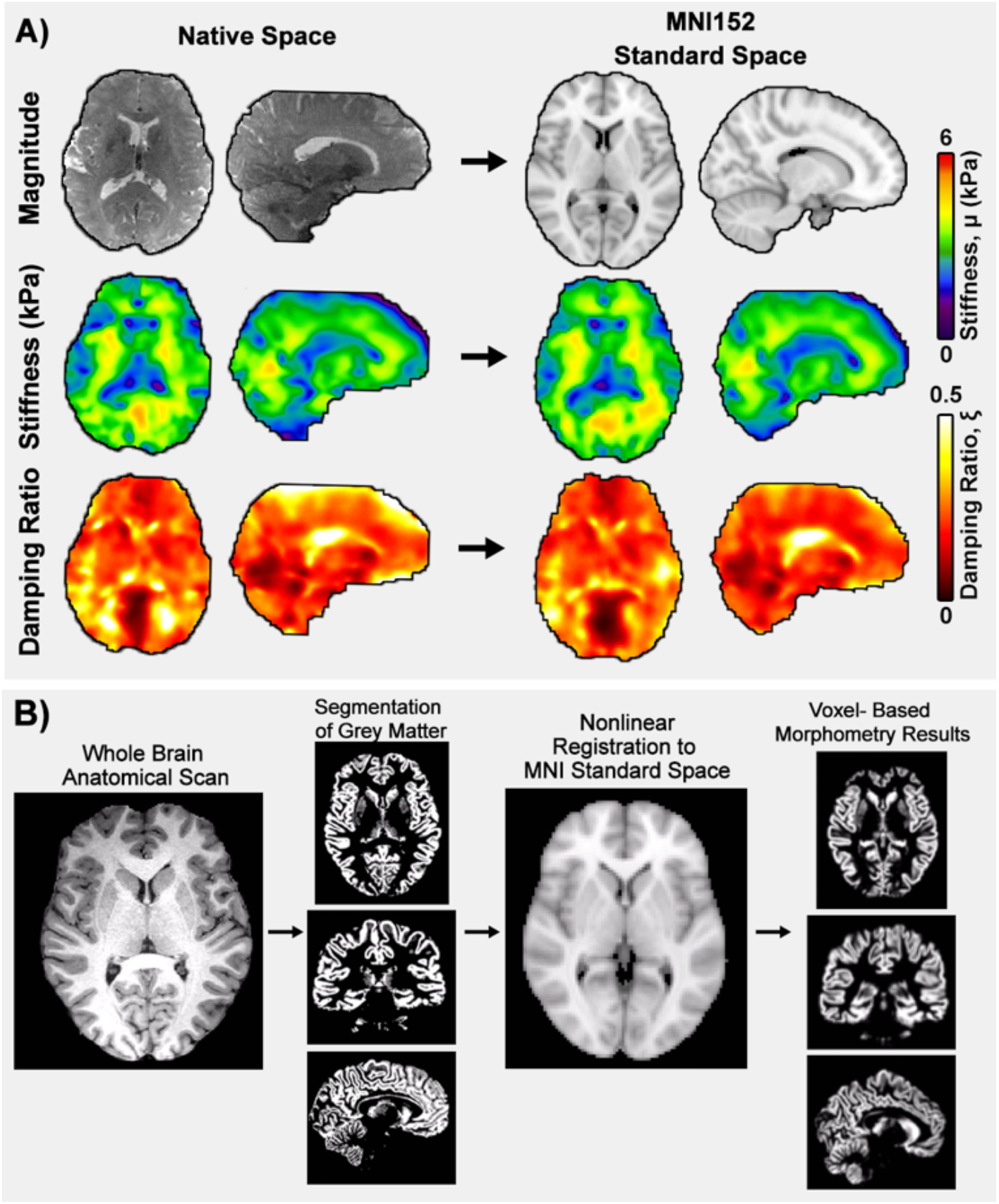
Overview of the major data processing steps. A) Example anatomical scan, stiffness map, and damping ratio map before and average conversion to MNI152 standard space. B) Steps for generating volume maps using voxel-based morphometry.

Volumetric maps were created using voxel-based morphometry (VBM) in FSL-VBM (Ashburner & Friston, 2000; S. M. Smith et al., 2004). Several previous studies have shown that VBM measures of brain volume are sensitive to age-related changes to the brain (C. D. Smith et al., 2007; Taki et al., 2004; Tisserand et al., 2004). In the VBM pipeline, the T1-weighted anatomical images were brain-extracted and gray matter was segmented using FSL FAST (Zhang et al., 2001). Data was then registered to the MNI152 standard space using the FNIRT non-linear registration. The resulting images are averaged and flipped along the x-axis to create a left-right symmetric, study-specific grey matter template. Then, all gray matter segmented images were non-linearly registered to this study-specific template. The gray matter images were then smoothed with an isotropic Gaussian kernel with a sigma of 8 mm. The resulting images depict the morphometry of gray matter regions relative to the rest of the cohort and are effectively voxel-wise maps of volumetric measures. An overview of this process is shown in Figure 2B.

Prior to being input to the CNN, all maps of volume and mechanical properties were restricted to the middle 55 slices of the MNI template to account for the top and bottom slices being outside the imaging field-of-view for some subjects. After truncation, each brain map then had dimensions of 91 × 109 × 55 and was standardized so that the mean of the voxels across the entire sample was 0 and the standard deviation was 1. This method involves subtracting the mean of the dataset from each voxel and then dividing it by the standard deviation of the dataset. Missing values in the brain maps associated with voxels outside of the brain tissue were set to 0.

### 2.3 Convolutional Neural Network Model

A 3D CNN was implemented using the Keras API with a TensorFlow backend (Abadi, 2016; Arnold, 2017). The CNN, shown in Figure 3, was designed to have sequentially connected blocks that process 3D patterns in the brain map. Each block is a sequence of three layers: two transposed 3D convolutional layers and equipped with ReLu activations and a max-pooling layer that is preceded by a batch normalization procedure. The kernel size of all convolutional layers was 3 × 3 × 3 and the pool size of all max-pooling layers was 2 × 2 × 2. For our implementation, we arranged 5 blocks sequentially with *n*_*chan*_ × 2^*i*−1^ filters, *i* = 1,2,3,4,5 per block and chose *n*_*chan*_ = 8. The output of the last block was reshaped into a dense layer and then followed by 2 dense layers composed of 640 and 100 nodes respectively, equipped with ReLu activations. Finally, the last dense layer was fully connected to a single node equipped with a linear activation, which is the brain age prediction. The Adam optimization algorithm was used to optimize the model by minimizing the mean squared error (MSE) and was chosen because it adapts the step size of the stochastic gradient descent for each parameter, which helps to speed up training and is robust to noisy data (Kingma & Lei Ba, 2015). The recommended default values were used for *β*_1_, the exponential decay rate for the first moment estimate, for *β*_2_, the exponential decay rate for the second moment estimate, and for *ε*, a constant for numerical stability (Kingma & Lei Ba, 2015). These values are 0.9, 0.999, and 10^−8^ for *β*_1_, *β*_2_, and *ε*, respectively. The CNN architecture is similar to the architecture described by Cole et al. (2017). The validation performance was assessed in terms of mean absolute error (MAE) of the age prediction. All processing was done on the Caviness high-performance computing cluster at the University of Delaware using a Tesla T4 GPU and Intel E5 family processor cores. The source code is publicly available at: github.com/mechneurolab/mechanical_brain_age.

**Figure 3.**
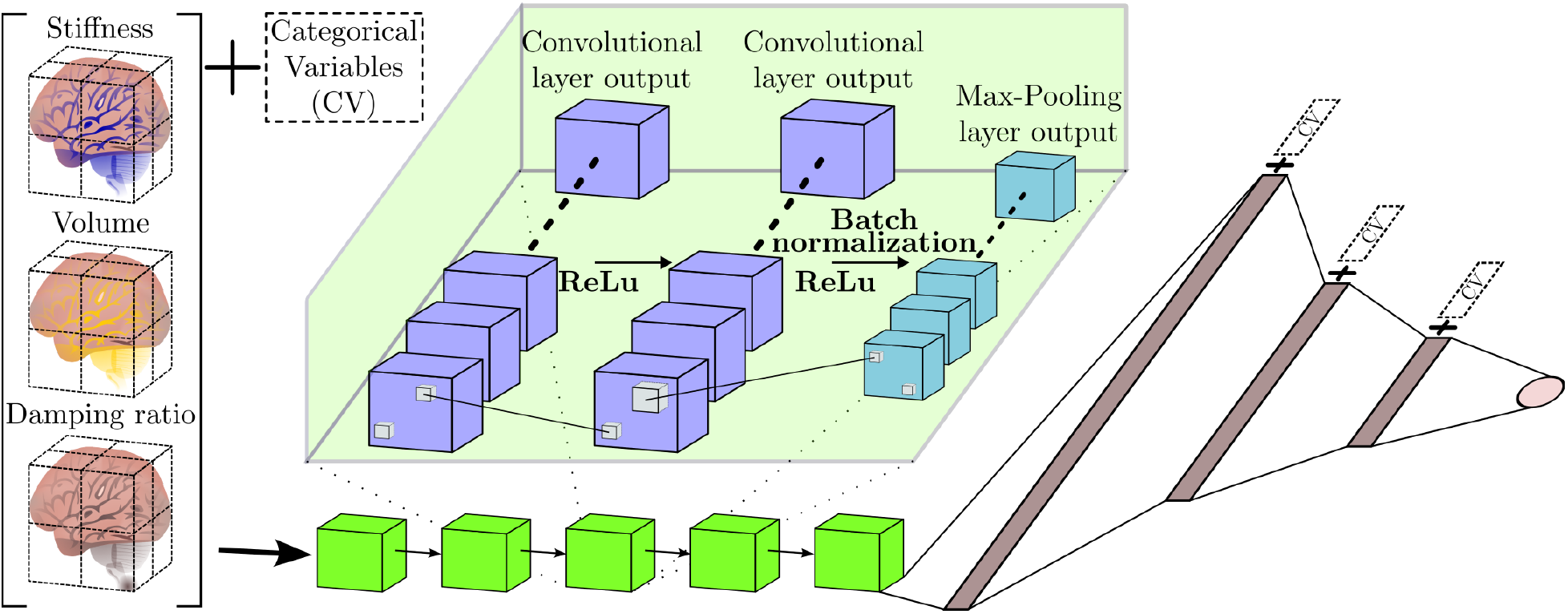
Overview of the CNN model architecture. As shown, the model is comprised of 5 blocks sequentially arranged, where each block is a combination of convolutional and max-pooling layers. The last layer of the fifth block is flattened and consecutively connected to two dense layers and a single node output. Combinations of categorical variables as inputs are evaluated at four different positions in the architecture. These combinations and positions are considered hyper-parameters.

From the dataset of 279 subjects, approximately 20% (57 subjects) were reserved for the test group and the remainder (222 subjects) were assigned to the training (166 subjects) and validation (56 subjects, 20% of the total dataset) groups. The dataset was sampled at random to assign subjects to these groups, and the random number generator in Python was seeded to ensure that the subjects assigned to each group remained consistent across models. Since all subjects were cognitively normal, their brain age was assumed to equal to their chronological age. To demonstrate the effectiveness of our CNN model relative to a traditional statistical model, we also used the training data to develop a basic regression equation to predict chronological age from average whole brain stiffness values. We applied this regression equation to the average stiffness values from the test set and computed the MAE of the predicted ages from the true chronological ages.

Stiffness is typically considered the primary outcome measure for MRE (Arani et al., 2021; Murphy et al., 2019), and has shown the strongest relationships with age (Hiscox et al., 2021), and as such was our primary brain structural measure of interest. To understand the effects of combining multiple physical property measures, we additionally tested CNN models that used five different combinations of imaging measurements: volume maps, damping ratio maps, stiffness and volume maps together, stiffness and damping ratio maps together, and a model with all three maps together. We determined the optimal hyperparameters for each of these models using Bayesian optimization with the validation set MAE as the cost.

### 2.4. CNN Hyperparameter Selection and Validation

We first tested an initial, unoptimized CNN model using a batch size of 12, a learning rate of 0.001, and 31 epochs. We did not include any categorical inputs in this model, and thus the architecture type was not meaningful. Then, to determine the best parameters for brain age prediction, we optimized the hyperparameters of the CNN model for different combinations of input maps using Bayesian optimization, which has been shown to be a fast and efficient method for hyperparameter tuning (Victoria & Maragatham, 2020). As shown in Table 2, we tested a variety of learning rates, batch sizes, architecture types, and categorical inputs. We also identified the optimal number of epochs using an upper bound of 60, since the validation set performance was used to determine the optimal number of epochs.

**Table 2.** Overview of the hyperparameters tested using Bayesian optimization

| Hyperparameter | Value |
| --- | --- |
| Learning Rate | [1e-5,1e-2] sampled logarithmically |
| Batch Size | 4, 8, 12, 16, 20, 24 |
| Architecture Type | '1','2','3','4' |
| Categorical Inputs | 'None', 'Sex', 'Study', 'Sex and Study' |
| Number of Epochs | ≤60 |

The Bayesian optimization procedure was executed for 60 trials, and the first 10 were performed with a random selection of hyperparameters. After the 10th trial, the optimization algorithm started picking the best combination of hyperparameters, considering the combinations that yielded the best results in the previous trials. The best hyperparameters were decided based on the mean absolute error of the validation set.

We tested the inclusion of additional categorical variables as inputs, including subject sex and the subject study ID, to the model to decrease the model bias. Subject sex was tested because, for at least some structures in the brain, aging effects may be more apparent in men than women (Coffey et al., 1998), and sexual dimorphism in brain mechanical properties has been reported previously (Arani et al., 2015; Hiscox, McGarry, et al., 2020; Sack et al., 2009). Since this project uses data collected from eight studies, a categorical variable corresponding to the particular study in which the subject participated was also added as an input to account for any differences resulting from combining data from multiple studies (Hiscox, McGarry, et al., 2020).

These categorical variables (sex and study identifier) were added to the model using Scikit-learn’s one-hot encoding function (Pedregosa et al., 2011), which maps each category feature to a binary vector. Concatenating the two one-hot vectors yielded a 10-dimensional vector (2 dimensions for sex and 8 for study ID). To compare models that ignore these categorical variables, the first two dimensions are set to 0 if sex is not being included as an input. As with sex, if study ID is not being included in the model, all 8 dimensions are set to zero.

We also defined four CNN architecture types to determine the best position to add the categorical vector to the image data, which are shown in Figure 3. The first design includes the categorical vector in the feature learning stage of the CNN so that each voxel was independently regressed on the categorical variable. In the second design, the categorical inputs are added to the first dense layer after the convolutional part of the neural network. The third architecture type attaches the categorical inputs to the nodes in the second dense layer after the convolutional segment. Finally, the fourth architecture type joins the categorical inputs with the nodes of the third layer after the convolutional section. As the architecture type number increases, the weighting that the model places on the categorical inputs decreases.

### 2.5. Model Performance Metrics

Model quality was primarily assessed using the mean absolute error (MAE) of the predicted age compared to the true chronological age on the test set. Root mean squared error, median absolute error, R^2^ values, and the average training and testing times were also computed and reported.

## 3. Results

### 3.1. Brain age prediction from stiffness maps

For the sake of comparison to our CNN model, we first used the training set to develop a linear regression model to predict age in years from global brain stiffness values. We applied this equation to the global stiffness values for each subject in the test set and found that our linear regression model was able to predict age with an MAE of 11.93 years. The individual predicted ages for each subject in the test set are shown in Figure 4A.

**Figure 4.**
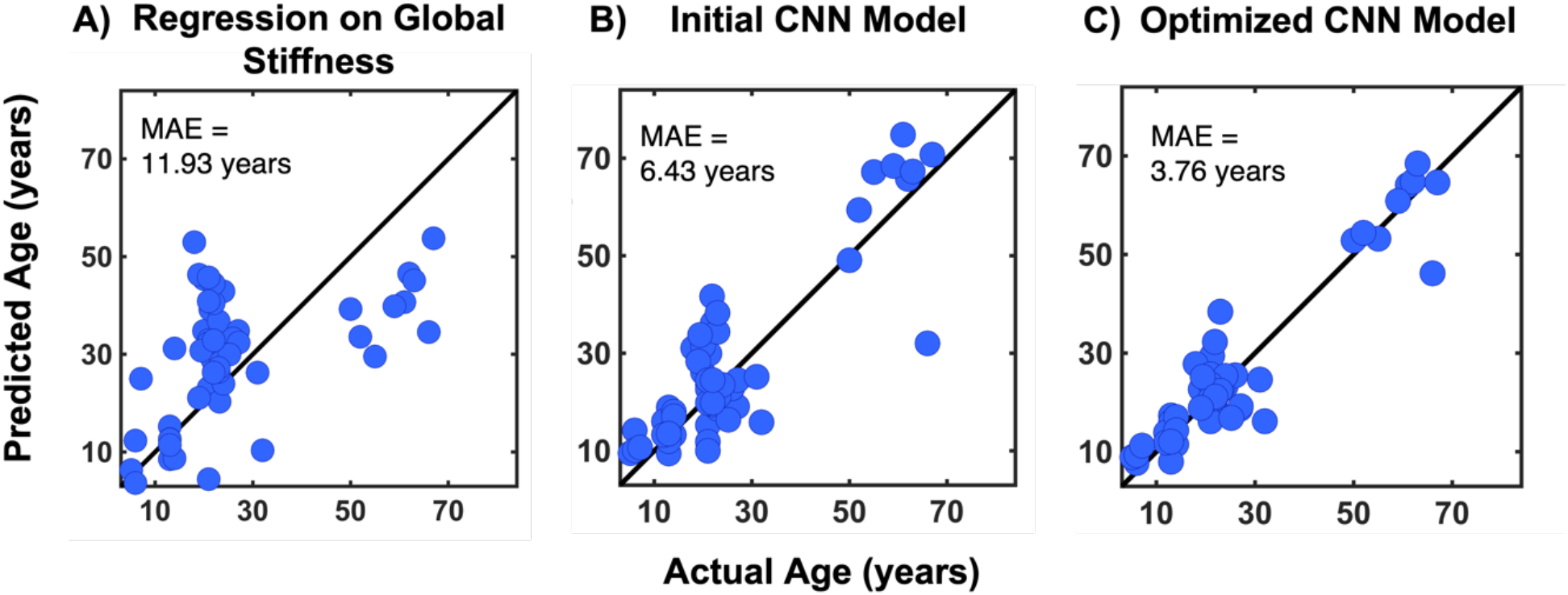
Results for the following models predicting age from brain stiffness: A) a linear regression model predicting age from global stiffness values, B) the initial CNN model with unoptimized parameters, and C) the optimized CNN model. Graphs are shown as actual by predicted age along with the unity line to illustrate perfect agreement.

We then used the CNN model to predict age from maps of brain stiffness using our initial, unoptimized hyperparameters and no additional categorical inputs (i.e. without sex and study ID), and these results are shown in Figure 4B. After training the CNN on stiffness maps from 222 individuals, we found that age can be accurately predicted with an MAE of 6.43 years. The hyperparameters corresponding to this model are a batch size of 12, a learning rate of 0.001, and 31 epochs. In general, the model performed better on the younger subset (ages 5–32) of individuals in the test group (MAE = 5.75 years) than the older subset (ages 50-67) of individuals in the test group (MAE = 9.95 years). These results far outperformed the basic regression we conducted using a single average value of whole brain stiffness to predict age.

To determine the best hyperparameters for predicting brain age from brain stiffness maps, we optimized the learning rate, batch size, architecture type, categorical inputs, and the number of epochs. We found that using no additional categorical inputs, a learning rate of 0.0002, a batch size of 4, and 50 epochs produced optimal results. Since no categorical inputs were being added, the architecture types were all the same, and the optimal architecture type was not meaningful. As shown in Figure 4C, this model had an MAE of 3.76 years. This model performed similarly for the younger set of individuals in the test group (MAE = 3.58 years) and the older subset of individuals in the test group (MAE = 4.71 years). This suggests that optimizing the hyperparameters helped the model to generate more accurate predictions across the entire age range. Training and testing times and the optimal hyperparameters for this model are shown in Table 3, and additional metrics for the performance of this model are shown in Table 4.

**Table 3.** Overview of average training and testing times for each model and optimal hyperparameters determined using Bayesian optimization

| Input | Average Training Time (minutes) | Average Testing Time (seconds) | Best Epoch | Categorical Inputs | Architecture Type | Learning Rate | Batch Size |
| --- | --- | --- | --- | --- | --- | --- | --- |
| Stiffness | 7.74 | 0.02 | 50 | None | N/A | 0.0002 | 4 |
| Volume | 7.52 | 0.02 | 47 | Sex, Study | 4 | 0.01 | 4 |
| Damping Ratio | 6.57 | 0.02 | 50 | Study | 1 | 0.01 | 16 |
| Stiffness, Volume | 7.12 | 0.02 | 51 | Study | 1 | 0.01 | 12 |
| Stiffness, Damping Ratio | 7.25 | 0.02 | 50 | Sex, Study | 1 | 0.002 | 16 |
| Stiffness, Volume, Damping Ratio | 7.75 | 0.02 | 37 | Sex, Study | 3 | 0.001 | 12 |

**Table 4.** Calculated mean absolute error, root mean square error, median absolute error, and R^2^ for each model

| Input | Mean Absolute Error (years) | Root Mean Square Error (years) | Median Absolute Error (years) | $R^2$ |
| --- | --- | --- | --- | --- |
| Stiffness | 3.76 | 5.52 | 2.64 | 0.88 |
| Volume | 4.60 | 6.76 | 2.83 | 0.82 |
| **Damping Ratio** | 3.82 | 7.84 | 1.67 | 0.76 |
| **Stiffness, Volume** | 3.60 | 6.67 | 1.97 | 0.82 |
| **Stiffness, Damping Ratio** | 5.49 | 9.45 | 3.25 | 0.64 |
| **Stiffness, Volume, Damping Ratio** | 5.27 | 8.68 | 3.53 | 0.70 |

### 3.2. Inclusion of additional brain structural measures in CNN

Next, we investigated whether including additional measures of brain tissue (volume and damping ratio) to the CNN model improved its accuracy in predicting age. We used Bayesian optimization to determine the optimal hyperparameters for models using different combinations of the measures shown in Table 2. We first tested models for volume alone and damping ratio alone; these results are found in Figure 5A. We found that volume alone produced an MAE of 4.60 years, which was higher than the error for stiffness of 3.76 years. Damping ratio alone produced an MAE of 3.82 years, which was also slightly worse than stiffness.

**Figure 5.**
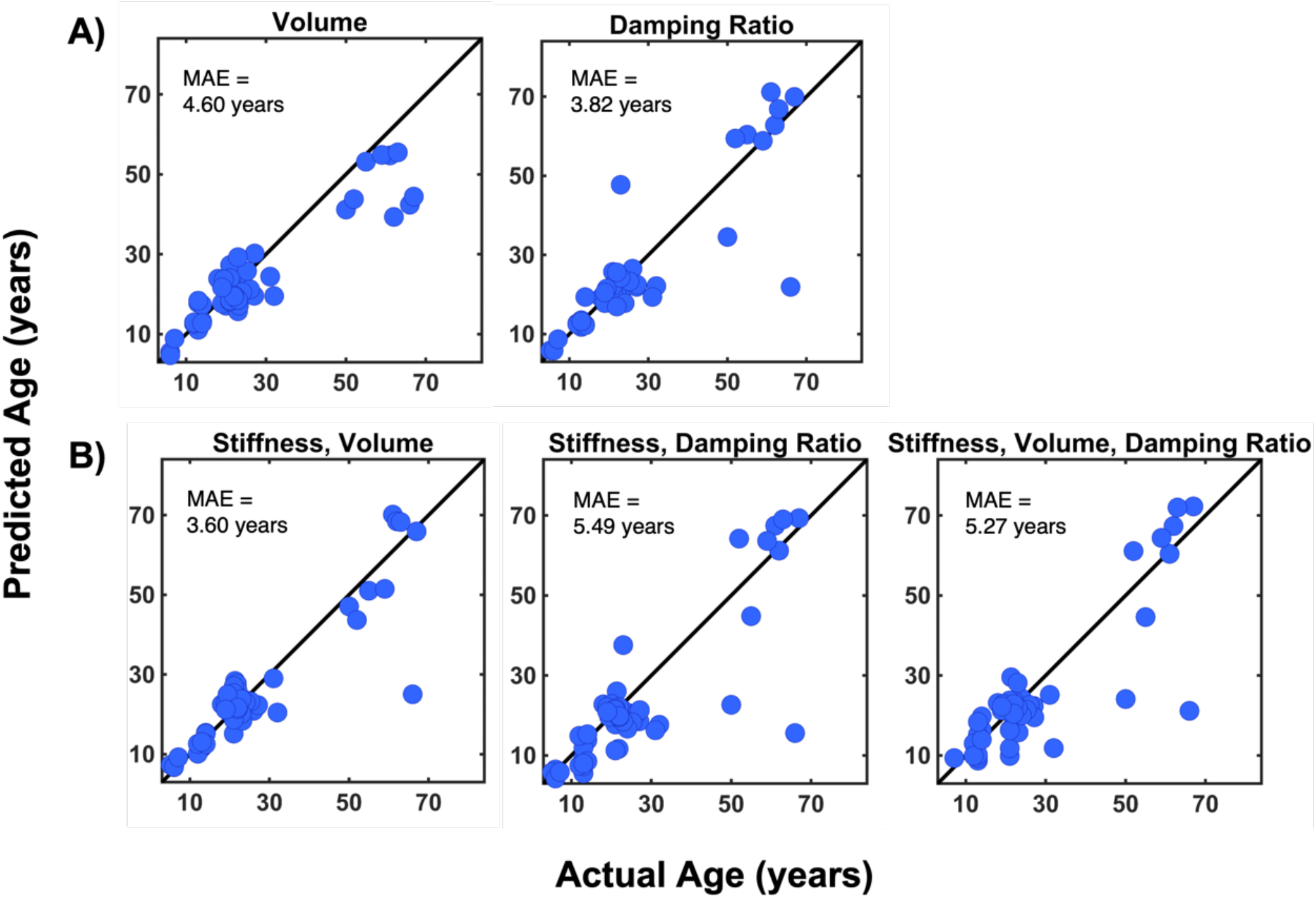
Results for each model with optimized hyperparameters including A) volume and damping ratio separately and B) combinations of brain maps including stiffness and volume; stiffness and damping ratio; and stiffness, volume, and damping ratio.

Figure 5B shows the effectiveness of the combinations of stiffness maps and the other brain structural maps. Stiffness and volume together produced an MAE of 3.60 years. This was a better predictor of brain age than stiffness maps alone (MAE: 3.76 years) and volume alone (MAE: 4.60 years). Including damping ratio along other measures, however, resulted in worse prediction performance. Stiffness and damping ratio maps together as model inputs produced an MAE of 5.49 years, while stiffness, damping ratio, and volume maps together produced an MAE of 5.27 years. Performance of both models was worse than models with each of the measures individually. A summary of the outcomes for each input combination is shown in Table 4.

## 4. Discussion

We have developed a machine learning methodology that yields a CNN model for predicting age from brain mechanical properties imaged using MRE. A CNN is advantageous over a traditional statistical model because it can analyze all brain regions simultaneously without explicit image segmentation and can learn to focus on the voxels in the brain map that accurately reflect the individual’s age. With a traditional statistical model, mechanical properties in a series of regions-of-interest from segmentation of brain regions need to be chosen and calculated as the set of variables for predicting brain age. This naturally reduces the dimensionality of the MRE data, losing the richness of the native images. Because stiffness is not homogeneous across brain regions (Guo et al., 2013), averaging stiffness values within a region can also lead to a loss of information. Further, using whole brain maps of mechanical properties, the CNN can learn inter-structure relationships and the patterns of how whole brain networks mature and degrade, without being restricted by pre-defined, regional boundaries.

For the first time, we have demonstrated that MRE measures can be used to successfully predict brain age. Our CNN model, which used maps of brain stiffness, predicted age with a mean absolute error of 3.76 years, which is comparable or even superior to previously published brain age models based on different neuroimaging data. This strong performance confirms that brain stiffness is sensitively affected by aging, likely by reflecting microstructural changes occurring throughout the lifespan. Previous MRE studies have pointed to these relationships with age (Arani et al., 2015; Delgorio et al., 2021; Sack et al., 2011), and here we advance those ROI-based analyses. The strong performance of the model to predict age, even in the absence of very large sample sizes for training, indicate that MRE outcomes should be considered in future brain age models.

To optimize our model, we used Bayesian optimization to determine the best model hyperparameters. Including sex as an input was expected to improve the model because existing research has shown that healthy females may tend to have more stiff brains than healthy males of the same age (Sack et al., 2009). However, for the stiffness model, the optimal hyperparameter combination did not include the sex variable as an input, indicating that the predictive value was either redundant or negligible, i.e. that the stiffness features most strongly reflective of age were not affected by sex. We also tested whether including the study identifier for each subject improved the model; while we did not expect significant differences between the studies, given very similar imaging and analysis protocols, we wanted to account for potential bias effects such as sequence type, resolution, and MRI scanner model. We found that the optimal hyperparameter combination for stiffness maps also did not include the study variable, indicating that the potential study effect on stiffness maps is minimal at best.

We sought to examine how brain stiffness performed in the CNN model compared to volume, a typical input to machine learning-based brain age models. Volume is a well-studied structural measure for predicting brain age and has previously been found to predict age with a high level of accuracy (Cole et al., 2017). We found that volume maps alone produced the highest error of any of the three brain structure measures, but that inputting volume and stiffness maps simultaneously performed slightly better than stiffness maps alone (MAE: 3.60 years and 3.76 years, respectively). This suggests that while stiffness appears to outperform volume alone in predicting age, both measures are complementary, providing unique information about brain age. These results are supported by a study which found that stiffness effects exist even when the volume of the region-of-interest is accounted for (Hiscox et al., 2018) and suggests that MRE measures provide additive value to volumetric measures.

We also examined the model that included maps of tissue damping ratio, the other common measure from MRE. Stiffness and damping ratio are independent measures, and it is expected that damping ratio reflects an organizational aspect of the tissue while stiffness reflects a compositional aspect of the tissue (Sack et al., 2013). The CNN model with damping ratio alone produced a slightly higher MAE than stiffness (MAE: 3.82 years and 3.76 years, respectively). However, including damping ratio together with stiffness in the CNN model performed worse than either measure separately (MAE: 5.49 years compared to 3.76 years or 3.82 years, respectively). Differences in brain damping ratio with age are less commonly reported (Hiscox et al., 2021), but have been observed in brain maturation (McIlvain et al., 2018) and in aging (Delgorio et al., 2021). Hiscox et al. reported a 21% difference in hippocampal damping ratio between younger and older adults (Hiscox et al., 2018), but other brain regions did not differ in damping ratio. Damping ratio has shown age effects, but these effects are weaker than those of stiffness. This may be because damping ratio reflects more individual differences in tissue integrity than are dictated by age. For instance, damping ratio of specific brain regions has been related to differences in cognitive function across several different types of behavioral assessments (Hiscox, Johnson, McGarry, et al., 2020; Johnson & Telzer, 2018; Schneider et al., 2022; Schwarb et al., 2016, 2019). Thus, the features of damping ratio maps distinct from stiffness are not solely influenced by age, leading to heightened error.

The specific mechanism which causes brain stiffness to be highly sensitive to age-related changes is an area of ongoing research. It is well established that large scale structural brain changes occur across the life span which impact measurable brain mechanical properties. During development, synaptic pruning of neural connections can cause a decline in gray matter volume (Gennatas et al., 2017), which occurs while myelination in the brain is increasing (Mabbott et al., 2006). Interestingly, we recently reported that brain stiffness appears to decrease from childhood to early adulthood (McIlvain, Schneider, et al., 2022). During normal adult aging, the brain undergoes a variety of degenerative changes, including decreases in the density of neurons, breakdown of myelin integrity, and the appearance of white matter lesions (Guttmann et al., 1998; Terry et al., 1987; Vernooij et al., 2008). With increasing age, the brain has been seen to become softer, with different regions of the brain having different rates of softening (Arani et al., 2015; Peters, 2006).

This softening can be exacerbated by neurological disorders. For instance, stiffness measures have been shown to be sensitive to demyelination caused by multiple sclerosis; one study found a 13% decrease in cerebral viscoelasticity in MS patients compared to healthy volunteers (Wuerfel et al., 2010). Stiffness has also been shown to be sensitive to changes caused by Parkinson’s disease, including a significant reduction in lentiform nucleus stiffness between PD patients and healthy, age-matched controls (Lipp et al., 2013). Alzheimer’s disease has been most commonly studied with MRE, and studies have shown lower stiffness in regions particularly affected by the disease (Gerischer et al., 2018; Hiscox, Johnson, McGarry, et al., 2020; Murphy et al., 2011, 2016). Recent studies have shown that using stiffness of particular regions can serve as effective biomarkers for differentiating between Alzheimer’s and other forms of dementia (ElSheikh et al., 2017; Pavuluri et al., 2023). This heightened softening in disease, particularly in patterns reflecting the distribution of pathology in the brain, point to the potential utility in a brain age model to identify and classify neurological conditions based on the difference with chronological age.

This study had several limitations. One limitation was that the data for the CNN was combined from multiple studies with slightly different imaging protocols and scanner hardware, although we accounted for this by including a study identifier as an input to the model. The results of our hyperparameter optimization also indicated that any study effect would be minimal. Some of the studies had different data resolutions, and even though it is all registered to a standard 2 mm resolution template, the underlying data remains different. However, previous studies have used data that were initially different resolutions and were registered to the same template and still demonstrated robust results (Hiscox, McGarry, et al., 2020). There was also uneven distributions of age or sex in some of the studies. Since the subject age distribution skewed towards younger adults and children, it may have been difficult for the model to learn the specific patterns of aging seen in older adults that are not strongly reflected in younger people. As the field of brain MRE expands, this CNN should be recalibrated with larger data sets to form even more robust predictions of brain age. Additional demographic inputs should also be included in the model to account for factors that may have positive or negative effects on brain tissue integrity, such as years of education or socioeconomic status. Finally, the CNN can be used to understand the features of stiffness maps that the model has identified as being most important for brain age prediction, which could provide information beyond what can be seen through traditional regional analysis about how the brain changes with aging.

## 5. Conclusion

This is the first study to use a CNN to predict brain age from MRE measures of brain stiffness. We found that MRE stiffness maps outperformed volumetric maps in determining brain age and that when inputted simultaneously, brain stiffness and volume give a higher prediction accuracy than either metric individually. The findings of this study suggest that MRE measures may be more sensitive to the maturation and tissue degradation process than other structural imaging techniques and provide a more accurate way to quantify atypical aging. This study provides a framework which could eventually be used for screening purposes in a clinical setting to identify individuals deviating from the healthy brain aging trajectory. Using machine learning models for brain age prediction is particularly valuable since they can detect patterns or abnormalities not able to be detected by the human eye, which is especially important for detecting minor changes before the onset of symptoms.

## 6. Acknowledgments

This work was supported in part by grants from the National Institutes of Health: R01-AG05885, R01-EB027577, U01-NS112120. This research was supported in part through the use of Information Technologies (IT) resources at the University of Delaware, specifically the high-performance computing resources.

